# A Fungal Spore Calendar for England: Analysis of 13 years of Daily Concentrations at Leicester, UK

**DOI:** 10.1101/2023.12.15.571848

**Authors:** F. A. Symon, S. Anees-Hill, J. Satchwell, A. Fairs, R. Edwards, A. J. Wardlaw, L. Cuthbertson, A. Hansell, C. H. Pashley

## Abstract

**Background:** Fungal respiratory allergy is believed to affect up to 30% of hayfever sufferers and up to 70% of severe asthmatics in the UK, however trends in fungal spore seasonality are not well described. Information about seasonal trends would help allergists determine sources of fungal sensitisation and aid disease management.

**Method:** Daily monitoring was carried out at Leicester from 2007 to 2020 using a Burkard volumetric spore trap. Fungal spore concentrations were analysed by microscopy, identifying 23 morphologically distinct taxa. Daily average concentrations were calculated as spores/m^3^ of air sampled and a 90% method used to determine the spore seasons.

**Results:** Thirteen years of data were used to develop a fungal spore calendar for the nine most abundant spore types identified; *Alternaria, Cladosporium, Didymella, Leptosphaeria, Sporobolomyces, Tilletiopsis* and *Ustilago* plus the wider groupings of *Aspergillus*/*Penicillium* type and coloured basidiospores. All have been implicated in fungal allergy.

We observed long seasons for, *Cladosporium, Sporobolomyces* and *Tilletiopsis*, beginning in late spring and ending in late autumn. In contrast *Ustilago* and the highly allergenic *Alternaria* showed relatively short seasons, spanning summer and early autumn. Temperature and precipitation were the main meteorological factors related to spore concentration with wind speed appearing to have little influence. Over the study period, there was a reducing trend for total spore concentrations, driven by a reduction in “wet weather” spores, in line with a reduction in precipitation. Conversely, the “dry weather” spores of *Alternaria* and *Cladosporium* demonstrated an increasing trend.

**Conclusion:** We present an aeroallergen calendar to provide readily accessible information to patients, healthcare professionals and pharmaceutical companies on exposure concentrations over the year in central England and potentially more widely across the UK. More research on allergenic thresholds would enhance the clinical usefulness of aeroallergen calendars.

## 1 Introduction

There are an estimated 2.2 – 3.8 million species of fungi, of which approx. 100,000 (or 2.6 - 4.5%) have been described (Hawksworth and Lücking, 2017). Fungi occupy a wide variety of environments and are important animal and plant pathogens, having a detrimental effect on agriculture and the health of both humans and animals. They reproduce by producing spores, many of which have adapted for airborne dispersal, and can exceed airborne pollen concentrations by 100 – 1000-fold in air samples (Horner et al., 1995). The ubiquitous distribution of fungi and the microscopic size of their spores means humans inhale large numbers of spores, alongside other airborne particles, such as pollen, dust, and air pollutants, on a daily basis.

The role of fungi as causative agents of allergy, asthma, non-invasive colonisation and opportunistic, life-threatening infections is becoming increasingly recognised (del Giacco et al., 2017; Knutsen et al., 2012; Rick et al., 2016). Fungi have been shown to play an important role in respiratory disease in both indoor and outdoor environments, with more than 112 fungi recognised as allergens (Levetin et al., 2016; Twaroch et al., 2015). An estimated 3-10% of the general population is thought to be sensitised to fungi and this increases to 12-42% of atopic patients and as much as 70% of patients with severe asthma may be sensitised to fungi, although estimates do vary depending on location and type of test used (Rick et al., 2016). There is a clear association between life-threatening asthma and sensitisation to fungal allergens, and studies have correlated outdoor spore concentrations with asthma symptoms (Batra et al., 2022; Newson et al., 2000; Pulimood et al., 2007; Tham et al., 2016). Fungal sensitisation may also pose a risk for the onset of asthma as has been observed in multiple studies (Denning et al., 2006; Vicencio et al., 2014; Zureik et al., 2002). Fungal spores have also been implicated in episodes of thunder storm asthma, with high levels of *Alternaria*, *Cladosporium, Didymella and* Sporobolomyces spores being associated with increased hospital admissions for asthma during or shortly after the storm (Allitt, 2000; Dales et al., 2003; Packe and Ayres, 1985; Pulimood et al., 2007).

Of key interest in fungal spore research is spore distribution and seasonal patterns, which can have important influences on health and can affect the agricultural environment and food production. Accurate information on fungal spore prevalence as well as forecasts can support improved allergy diagnosis, allergen avoidance, and symptom management, improving quality of life.

While much research has focused on the creation of pollen seasonal calendars to aid allergen avoidance, the same cannot be said in relation to annual fungal spore distributions. Studies that put forward fungal spore calendars specifically have tended to study just one or two spore genera or phyla (Antón et al., 2019; Elvira-Rendueles et al., 2019; Sánchez Reyes et al., 2022) or to sample for less than the 5 years recommended by the European Aerobiological Society (Antón et al., 2019; Bednarz and Pawłowska, 2016; Galan et al., 2017; Kasprzyk et al., 2004; Martínez-Bracero et al., 2022; Reyes et al., 2016; Scevkova and Kovac, 2019).

Using one of the longest sampling studies of airborne fungal spores to date, this publication aimed to explore fungal spore trends over a 13-year period to provide a fungal spore calendar for the UK. We describe the fungal content of Leicester’s atmosphere from 2007-2020 for the most abundant spore types seen in Leicester, central England, relating the distribution of spores to meteorological parameters in order to describe trends in spore data that may have impacts on the diagnosis, avoidance and management of fungal allergy sufferers.

## 2 Materials and Methods

### 2.1 Site location and air sample collection

The city of Leicester (52°38’ N 1°5’ W) is located in Leicestershire, in the East Midlands of England, UK. Leicester is an industrial city with a population of approximately 368,600 in 2021 (Office for National Statistics, 2022, June 28) and is located about 90km from the coast. The city is predominantly surrounded by agricultural land (Pashley et al., 2009).

Sampling of airborne fungal spores was conducted continuously from 1^st^ March to 30^th^ November, spanning the years 2007 to 2020. Missing dates unavailable for analysis are recorded in Table S1. We excluded 2016 due to insufficient data. A 7-day recording volumetric spore trap (Burkard Manufacturing Co. Ltd.) was used to sample the air and was situated 12m above ground level (60m above sea level) on the roof of the Bennett building on the University of Leicester campus. This particular area of the city is characteristically urban, approximately 1km south of the city centre and next to 69 acres of open parkland with tree lined avenues.

Sampling slides were prepared according to standard procedures used for over 50 years in the UK and other European countries (Bednarz and Pawłowska, 2016; Newson et al., 2000; Sadys et al., 2016a; Skjoth et al., 2016). Twenty-three fungal spore types were identified by light microscopy (Eclipse Ci, Nikon or AxioScope A1, Carl Zeiss), 18 to genus level. Some fungal spores were counted jointly or as groups because visual distinction was not possible between genera, as in the case of *Aspergillus* sp. and *Penicillium* sp., or using groupings based on similar morphology, as for ascospores, hyaline basidiospores, coloured basidiospores and “rusts and smuts”. Spores were counted along one central transverse under x600 or x630 magnification, before applying a correction factor to give an estimation of the total number of spores per cubic metre of air per 24 hours (spores/m^3^).

### 2.2 Weather data

Minimum and maximum temperature, precipitation, wind speed and direction were provided by Leicester city council air quality group. The meteorological station was located 5km from the trap site. The hourly data received was independently averaged to provide the daily weather data or summed in the case of precipitation to give total daily precipitation. Directional wind averages were calculated using circular means.

### 2.3 Analysis

Annual cumulative concentrations were calculated from the summation of daily average spore concentrations from 1^st^ March to 30^th^ November of each year for the 23 morphologically distinct taxa or groups identified at this location (Table 1). Here we present the eight most abundant fungal taxa, identified to genus level, plus coloured basidiospores (highlighted in Table 1). The groupings of ascospores and hyaline basidiospores, although highly abundant, were considered too heterogeneous for further analysis. All of the 9 spore types analysed have been implicated in fungal allergy.

**Table 1.** Characteristics of fungal spore distribution for all spore types identified. Fungal spores were categorised to genus level when applicable, otherwise spores were grouped by visual distinction. Mean counts were calculated yearly from March to November and averaged across the 13 years of data collection. Maximum daily peak was calculated yearly as the highest number of spores recorded in a day for each spore type, during the recording period (March to November). Spore types in bold and indicated with * were included in further analysis, as they exceeded a mean daily maximum peak of 1000 spores/ m^3^.

| Spore Type | Mean yearly<br>spore<br>concentration<br>[March - Nov]<br>(spores/m<sup>3</sup>) | Mean yearly<br>spore<br>concentration<br>[March -<br>Nov] SD | % Total<br>spore<br>concentration | Mean Daily<br>Maximum<br>Peak<br>(spores/m<sup>3</sup>) | Greatest<br>Maximum<br>Daily Peak<br>(spores/m<sup>3</sup>) | Smallest<br>Maximum<br>Daily Peak<br>(spores/m<sup>3</sup>) |
| --- | --- | --- | --- | --- | --- | --- |
| ***Cladosporium*\*** | 875,395 | 236,641 | 26.23 | 38,164 | 70,029 | 18,042 |
| Ascospores | 620,921 | 228,623 | 18.61 | 19,986 | 40,047 | 8,333 |
| ***Sporobolomyces*\*** | 485,845 | 260,066 | 14.56 | 50,388 | 139,446 | 8,382 |
| Hyaline basidiospores | 422,520 | 189,502 | 12.66 | 12,200 | 33,757 | 4,186 |
| *Tilletiopsis* * | 312,306 | 315,565 | 9.36 | 30,124 | 89,274 | 1,362 |
| Coloured basidiospores* | 136,452 | 32,187 | 4.09 | 3,967 | 8,455 | 2,654 |
| *Didymella* * | 104,532 | 109,978 | 3.13 | 8,035 | 27,342 | 967 |
| *Leptosphaeria* * | 36,495 | 18,193 | 1.09 | 1,426 | 2,440 | 465 |
| *Aspergillus/Penicillium*<br>type* | 34,715 | 9,754 | 1.04 | 1,272 | 2,715 | 496 |
| *Ustilago* * | 32,008 | 10,325 | 0.96 | 4,165 | 8,375 | 1,776 |
| *Alternaria* * | 25,353 | 10,258 | 0.76 | 1,655 | 3,700 | 918 |
| *Ganoderma* | 13,920 | 3,995 | 0.42 | 307 | 420 | 161 |
| Rusts/smuts | 13,870 | 6,197 | 0.42 | 448 | 724 | 220 |
| *Botrytis* | 12,797 | 5,242 | 0.38 | 460 | 1,190 | 248 |
| *Entomophthora* | 5,287 | 2,114 | 0.16 | 200 | 372 | 99 |
| *Epicoccum* | 3,876 | 1,145 | 0.12 | 186 | 397 | 79 |
| *Erysiphe* | 3,604 | 1,487 | 0.11 | 117 | 180 | 67 |
| *Polythrincium* | 2,499 | 1,241 | 0.07 | 135 | 337 | 30 |
| *Lewia/Pleospora* | 2,492 | 976 | 0.07 | 173 | 459 | 43 |
| *Torula* | 1,590 | 414 | 0.05 | 81 | 161 | 43 |
| *Stemphylium* | 1,282 | 1,459 | 0.04 | 191 | 620 | 24 |
| *Drechslera* | 353 | 193 | 0.01 | 43 | 140 | 12 |
| *Pithomyces* | 247 | 165 | 0.01 | 30 | 50 | 12 |

The 90% method (Skjoth et al., 2016) was used to define the spore season retrospectively with the start of the season being the date at which 5% of the total concentration for the 9 months was recorded and 95% being the end date of the season. The Seasonal Spore Integral (SSIn), which is the sum of the daily spore concentrations recorded during the defined spore season was then calculated. Allergenic thresholds for *Alternaria* sp. and *Cladosporium* sp. were defined as 100 and 3000 spores per cubic metre respectively (Bagni et al., 1977; Gravesen, 1979; Rapiejko et al., 2007).

A Spearman’s rank correlation test was used to examine the relationship between daily spore concentrations and selected meteorological parameters within fungal spore seasons for each taxon investigated. Further analysis with multiple regression incorporating lags of up to 3 days for meteorological variables was carried out. Meteorological variable selection per model was performed using best subset selection with cross-validation, rather than metrics that penalise upon the addition of further variables. Spore counts were transformed using a logarithmic transformation (+1) as linear regression residuals were non-normal. Missing counts were imputed using interpolation from periods up to 7 days pre and post missing values, or the mean of surrounding years. Meteorological variables in the model included daily maximum air temperature (Tm, °C), wind speed (WS, m/s), sum of precipitation (P, mm), and mean circular wind direction (WD, degrees). Wind direction was discretised into 4 categories (Q1: north east, Q2: south east, Q3: south west, Q4: north west). For each taxon run, the model providing the lowest root mean square error (RMSE) in prediction using k-fold cross-validation (k = 10) was selected as the final model per fungal spore group. Long-term trends were also explored using simple linear regression across the overall 13-year SSIn. An alpha value of 0.05 was used as the threshold for statistical significance.

All analyses were performed using GraphPad Prism version 7.05.237 for Windows (GraphPad Software, San Diego, Ca), Microsoft Excel 365, and R version 4. 1. 2.

## 3 Results

### 3.1 Overall fungal spore contribution and distribution

The 23 fungal spore types routinely identified and analysed in air samples collected in Leicester during the 13-year period from 2007 to 2020 (excluding 2016) and their mean annual abundance from March - November are shown in Table 1. The most abundant spore type observed was *Cladosporium*, giving the highest average annual spore concentration of 875,395 spores/m^3^ and the highest spore concentration in 8 of the 13 years studied.

Ascospores formed the next most abundant group having the highest annual spore concentrations in 2017 and 2019, followed by *Sporobolomyces* (highest in 2010), hyaline basidiospores and then *Tilletiopsis* (highest in 2007 and 2012). Together these spore types accounted for 81.4% of the total spores observed.

Of the spores identified to fungal genus, *Cladosporium*, *Sporobolomyces*, *Tilletiopsis*, *Didymella*, *Leptosphaeria*, *Aspergillus*/*Penicillium* type, *Ustilago* and *Alternaria* exceeded a mean daily maximum peak of 1000 spores m^-3^ (Table 1). The remaining taxa occurred at much lower concentrations of less than 500 spores/m^3^ mean maximal daily peak and less than 15,000 spores/m^3^ mean yearly concentration. All spore types showed a large degree of variability in both their total (March – November) spore concentration and daily maximal peak concentration across the 13 years studied (Table S2). However, despite the variability in spore concentration, *Alternaria, Cladosporium*, *Didymella*, *Sporobolomyces*, *Tilletiopsis*, *Ustilago* and the coloured basidiospores showed a consistent pattern of distribution across the months in all 13 years studied (Figure 1), while, *Leptosphaeria*, and *Aspergillus*/*Penicillium* type were more variable.

**Figure 1:**
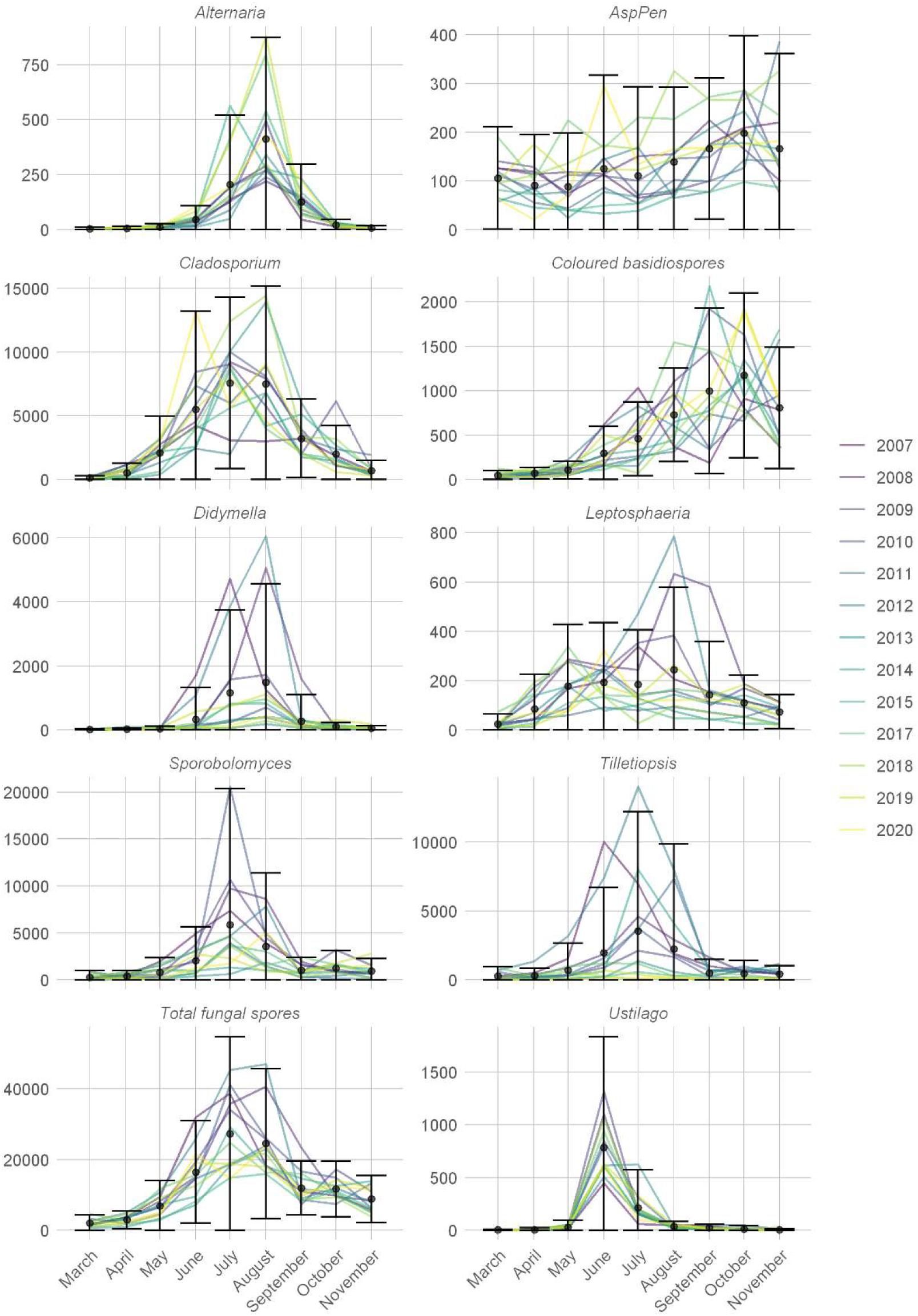
Annual distribution of spores over the 13 years sampled. Points display monthly means (spores/m^3^) with error bars displaying the standard deviation. Each year is represented by a single line.

*Ustilago* was the most consistent of the spores studied, peaking in June in 12 of 13 years, while *Alternaria* and *Didymella* concentrations increased through July, peaking in August. *Cladosporium*, *Sporobolomyces* and *Tilletiopsis* demonstrated a more prolonged peak from June to August. *Leptosphaeria*, *Aspergillus*/*Penicillium* type and coloured basidiospores demonstrated a more complex distribution with coloured basidiospores and *Aspergillus*/*Penicillium* type spores tending to appear in higher numbers towards the end of the summer and into the autumn months when other spores were declining in numbers. It should be noted that *Aspergillus*/*Penicillium* type and coloured basidiospores are groupings which include a very large number of species and by grouping them into single taxa the signal from subgroups may be lost overall.

### 3.2 Fungal spore seasons

The main spore season (MSS) was defined by the 90% method and is shown in the fungal spore calendar in Figure 2. The summer months of June to August were found to have the greatest diversity of fungal spore types, and 6 of the studied spores were found to peak during this period.

**Figure 2:**
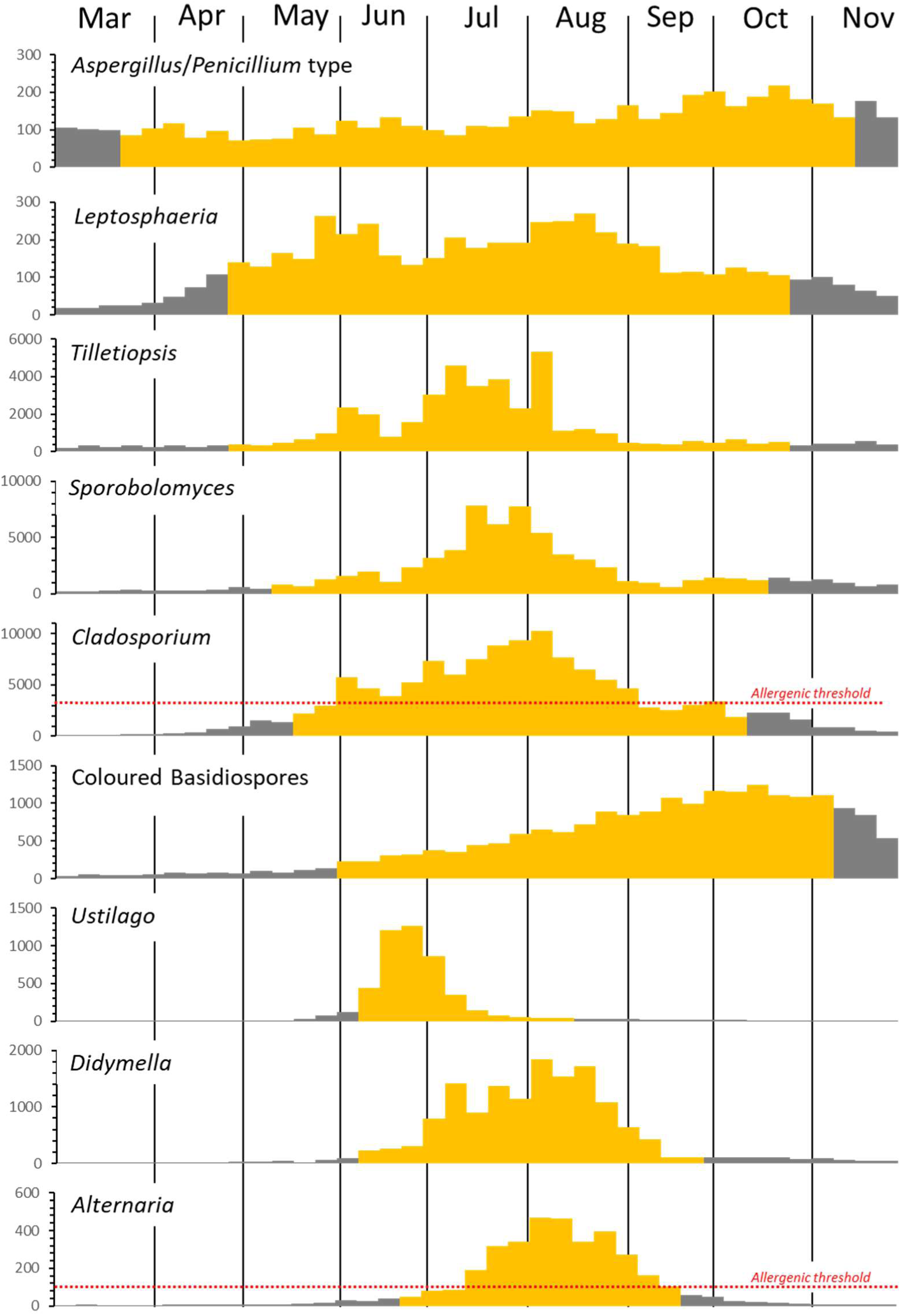
Fungal spore calendar for the 9 main spore types observed in Leicester. Spore types were ordered by the timing of their appearance. The main spore season (MSS) for each of the spores studied is shown in yellow (defined by the 90% method and based on 13 years of data). Allergenic threshold concentrations for *Alternaria* and *Cladosporium* are indicated by the red dotted line. No allergenic threshold is currently available for any other spore type.

*Aspergillus*/*Penicillium* type and *Leptosphaeria* had both the earliest and longest spore seasons, beginning in March and April. However, they had the lowest mean peak daily concentration and a relatively low SSIn (Table S2) indicating a prolonged low-level presence. In the case of *Leptosphaeria* no peak period could be defined with peak daily concentrations being scattered throughout the season.

Late April and May were found to be the beginning of the season for *Tilletiopsis*, *Sporobolomyces* and *Cladosporium*. Like *Aspergillus*/*Penicillium* type and *Leptosphaeria,* these spores had a relatively long season, continuing into October, but they demonstrated at least a 10-fold higher SSIn (Table S2) and had a clear peak period from late June until August, with maximal daily counts peaking in July, and into August for *Cladosporium* (Figure 1).

*Ustilago, Didymella* and *Alternaria* had the shortest seasons of the spores analysed, beginning in June, and ending in September, or August for *Ustilago*. These spores had similar SSIn to the earliest spores, however, appearing over a shorter period of time, they had a higher mean peak daily concentration (Table S2). They all had very distinct, short, peak periods, in June for *Ustilago*, and August for *Alternaria* and *Didymella* (Figure 1).

Coloured basidiospores and *Aspergillus*/*Penicillium* type spores tended to show increasing daily concentrations towards the end of summer and into the autumn months with peak periods from the end of August until November, reaching a maximum in October.

Although several of the fungal taxa discussed here are known to cause IgE sensitisation (Simon-Nobbe et al., 2008), allergenic thresholds are only available for *Cladosporium* (> 3000 spores/m^3^ per day) and *Alternaria* (> 100 spores/m^3^ per day) (Bagni et al., 1977; Gravesen, 1979; Rapiejko et al., 2007). Over the study period *Cladosporium* spore levels were found to have exceeded allergenic threshold on average on 88 days of the year, ranging from 52 days in 2012, which also recorded the lowest SSIn, to 120 days the year before (Table S2). Although 2011 had the longest *Cladosporium* season, and gave the highest number of days above allergenic threshold, this did not correspond to the highest SSIn, this was observed in 2018 (1,203,640 spores/m^3^, Table S2). 2018 had the second highest number of days above allergenic threshold (117 days). No correlation was observed between season length and the number of days above allergenic threshold.

The highly allergenic *Alternaria* was found to exceed published allergenic levels on average on 52 days, ranging from 33 days in 2012, also the year with the lowest SSIn, to 72 days in 2014. Although 2014 had the highest number of days above allergenic threshold for *Alternaria,* it did not have the highest SSIn, it was also the only year in which *Alternaria* spore concentrations peaked in July, rather than August, and carried on at significant levels into September (Figure 1). The highest SSIn for *Alternaria* was found in 2019 at 42,670 spores/m^3^, the year with the shortest season duration (72 days). Like *Cladosporium*, *Alternaria* showed no significant correlation between season length and the number of days above allergenic threshold.

*Alternaria* and *Cladosporium* have overlapping seasons (Figure 2) and it was found that in 5 of the 13 years studied, there was substantial overlap of days when both *Alternaria* and *Cladosporium* exceeded their allergenic levels (Figure 3). In a further 3 years >80% of the *Alternaria* allergenic days corresponded to *Cladosporium* allergenic days.

**Figure 3:**
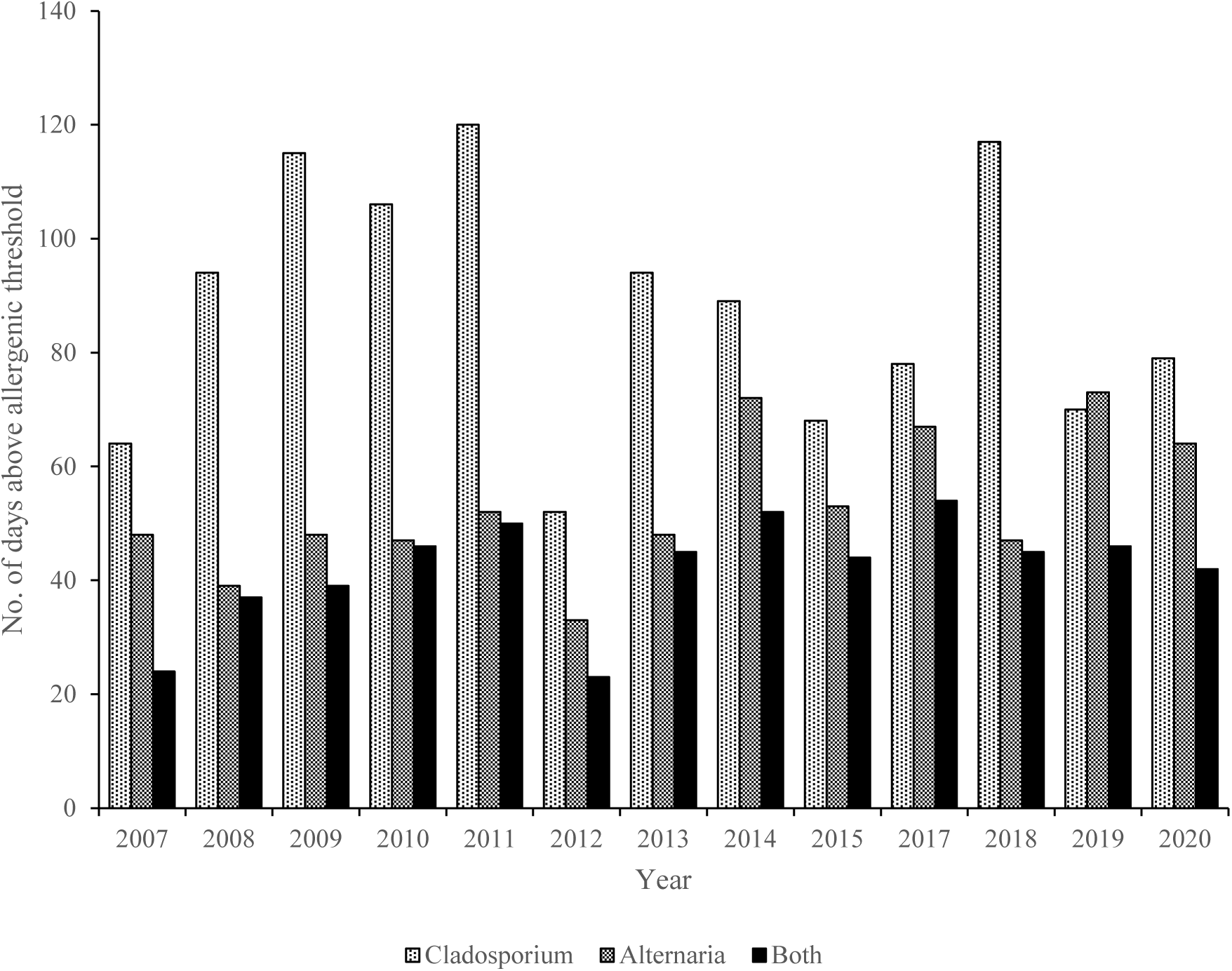
Number of days within each year when allergenic thresholds were exceeded during the study period. Allergic threshold is defined as; *Cladosporium* > 3000 spores/m^3^/day, *Alternaria* > 100 spores/m^3^/day. Stippled bars represent number of days *Cladosporium* > 3000 spores/m^3^/day, hatched bars represent number of days *Alternaria* > 100 spores/m^3^/day, solid bars represent number days when both taxa exceeded threshold on the same day.

### 3.3 Spore Dependence on Meteorological Parameters

Multiple linear regression was used to investigate potential associations between daily spore concentrations and meteorological parameters. The best set of variables and their corresponding lags were chosen via best subset selection. Using explained variance (R^2^) as an indicator of model fit, the models performed particularly well for *Cladosporium* and *Alternaria*, with the amount of explained variance being 59% and 46% respectively. Maximum daily temperature had a positive influence on *Cladosporium* and *Alternaria* spore concentrations, while wind speed on the previous day (lag1) had a negative influence (*Alternaria*; r = 0.154, p <0.01, *Cladosporium*; r = 0.159, p = <0.01). Total daily precipitation had a negative influence on *Alternaria* spore concentrations (r = −0.049, p <0.01), in agreement with the correlation analysis looking at each weather parameter in isolation (Spearman’s rank correlation, Table S3). In contrast, daily precipitation (P) on the previous day (lag1) had a positive influence on *Cladosporium* spore concentrations (r = 0.029, p <0.01). Wind direction also influenced *Cladosporium* spore concentrations, having a positive effect when coming from a south easterly (Q2) direction (r = 0.332, p <0.01) and a negative effect when coming from a south or north westerly direction (Q3; r =-0.153, p <0.01, Q4; r = −0.204, p < 0.01). A south-westerly wind also had a negative impact on *Alternaria* spore concentrations (r = −0.279, p <0.01).

The Multiple linear regression analysis (Table 2) demonstrated that meteorological parameters accounted for 14% – 18% of spore concentration variance for *Didymella*, *Sporobolomyces, Tilletiopsis* and *Leptosphaeria* with total daily precipitation having a significant positive effect on spore concentrations (*Didymella*; r = 0.093, p < 0.01, *Sporobolomyces*; r = 0.037, p < 0.01, *Tilletiopsis*; r = 0.066, p < 0.01 and *Leptosphaeria*; r = 0.038, p <0.01), in accordance with the analysis of each weather parameter individually (Table S3). A positive relationship with maximum daily temperature was observed for *Sporobolomyces* (r = 0.061, p <0.01) and *Tilletiopsis* (r = 0.048, p <0.01) however, maximal daily temperatures on preceding days (lags 1-3, Table 2) had as much or more influence on the spore concentrations of all four genera, *Didymella*, *Sporobolomyces, Tilletiopsis* and *Leptosphaeria*.

**Table 2.**
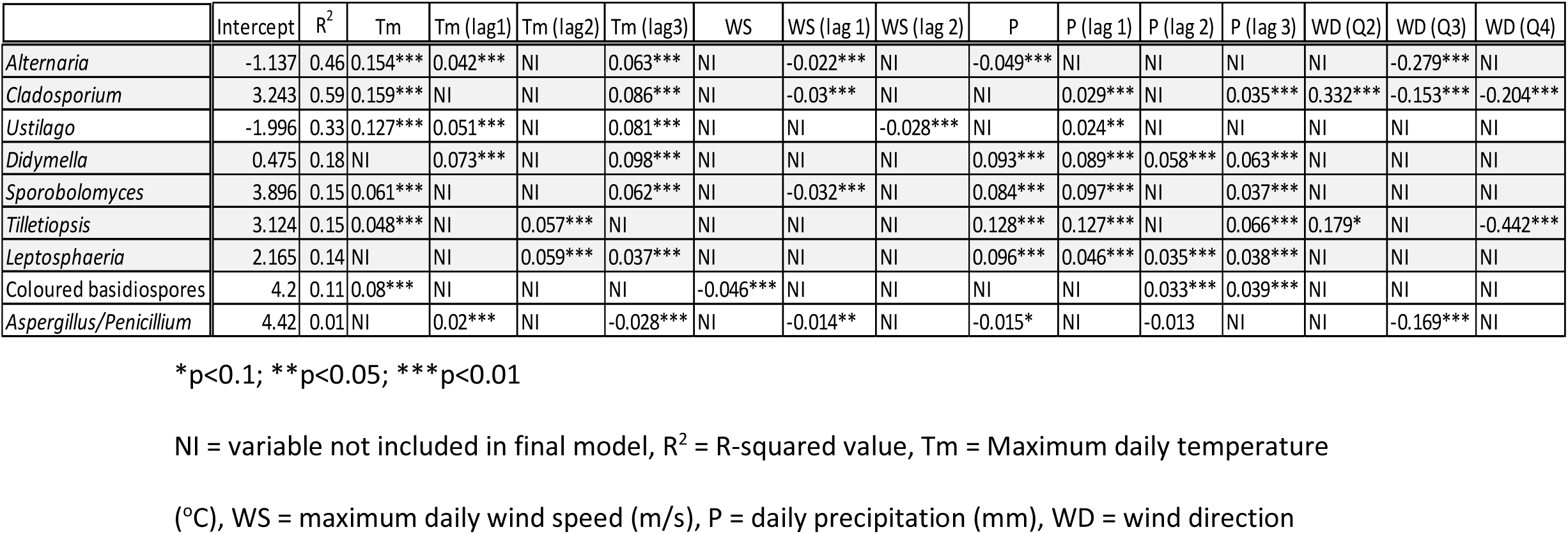
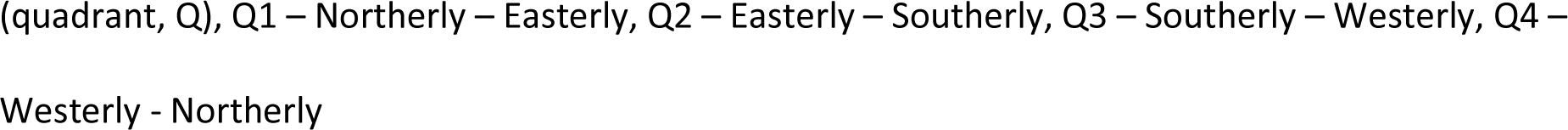
Multiple regression analysis of the effects of meteorological variables on various fungal spore concentrations.

|  | Intercept | R<sup>2</sup> | Tm | Tm (lag1) | Tm (lag2) | Tm (lag3) | WS | WS (lag 1) | WS (lag 2) | P | P (lag 1) | P (lag 2) | P (lag 3) | WD (Q2) | WD (Q3) | WD (Q4) |
| --- | --- | --- | --- | --- | --- | --- | --- | --- | --- | --- | --- | --- | --- | --- | --- | --- |
| *Alternaria* | -1.137 | 0.46 | 0.154*** | 0.042*** | NI | 0.063*** | NI | -0.022*** | NI | -0.049*** | NI | NI | NI | NI | -0.279*** | NI |
| *Cladosporium* | 3.243 | 0.59 | 0.159*** | NI | NI | 0.086*** | NI | -0.03*** | NI | NI | 0.029*** | NI | 0.035*** | 0.332*** | -0.153*** | -0.204*** |
| *Ustilago* | -1.996 | 0.33 | 0.127*** | 0.051*** | NI | 0.081*** | NI | NI | -0.028*** | NI | 0.024** | NI | NI | NI | NI | NI |
| *Didymella* | 0.475 | 0.18 | NI | 0.073*** | NI | 0.098*** | NI | NI | NI | 0.093*** | 0.089*** | 0.058*** | 0.063*** | NI | NI | NI |
| *Sporobolomyces* | 3.896 | 0.15 | 0.061*** | NI | NI | 0.062*** | NI | -0.032*** | NI | 0.084*** | 0.097*** | NI | 0.037*** | NI | NI | NI |
| *Tilletiopsis* | 3.124 | 0.15 | 0.048*** | NI | 0.057*** | NI | NI | NI | NI | 0.128*** | 0.127*** | NI | 0.066*** | 0.179* | NI | -0.442*** |
| *Leptosphaeria* | 2.165 | 0.14 | NI | NI | 0.059*** | 0.037*** | NI | NI | NI | 0.096*** | 0.046*** | 0.035*** | 0.038*** | NI | NI | NI |
| Coloured basidiospores | 4.2 | 0.11 | 0.08*** | NI | NI | NI | -0.046*** | NI | NI | NI | NI | 0.033*** | 0.039*** | NI | NI | NI |
| *Aspergillus/Penicillium* | 4.42 | 0.01 | NI | 0.02*** | NI | -0.028*** | NI | -0.014** | NI | -0.015* | NI | -0.013 | NI | NI | -0.169*** | NI |

\* $p < 0.1$ ; \*\* $p < 0.05$ ; \*\*\* $p < 0.01$
NI = variable not included in final model, R<sup>2</sup> = R-squared value, Tm = Maximum daily temperature (°C), WS = maximum daily wind speed (m/s), P = daily precipitation (mm), WD = wind direction
(quadrant, Q), Q1 – Northerly – Easterly, Q2 – Easterly – Southerly, Q3 – Southerly – Westerly, Q4 – Westerly - Northerly

Multiple parameter analysis suggested that 33% of *Ustilago* spore concentration variance was accounted for by meteorological parameters with maximum daily temperature (r = 0.127, p<0.01) and total daily precipitation on the previous day (lag1, r = 0.024, p = 0.05) having a positive influence, whilst wind speed two days beforehand had a negative effect (lag2, r = −0.028, p <0.01) (Table 2). Coloured basidiospores showed only 11% of spore variance was accounted for by weather parameters with maximum daily temperature having a positive influence (r = 0.08, p <0.01) as well as precipitation in the previous 2 – 3 days (lag 2; r = 0.033, p <0.01, lag 3; r = 0.038, p<0.01), while wind speed had a negative influence (r = −0.046, p<0.01) (Table 2).

### 3.4 Changes in seasonal fungal spore concentration

Linear regression revealed a statistically significant decrease in total fungal spore concentrations across the 13 years (β = −105,641, R^2^ = 0.36, p = 0.03), as did *Sporobolomyces* (β = −32,819, R^2^ = 0. 36, p = 0.03) and *Tilletiopsis* (β = −37,315, R^2^ = 0.31, p = 0.05), while *Alternaria* showed a significant increase (β = 1352, R^2^ = 0. 39, p = 0.02) (Table S4). The trend for *Didymella* and *Leptosphaeria* was decreasing, while *Cladosporium* showed an increasing trend (β = 5751, R^2^ = 0.013, p = ns). *Ustilago*, *Aspergillus*/*Penicillium* type and coloured basidiospores showed little change in SSIn across the 13 years of this study.

These trends remained the same when total spore concentrations from March – November were analysed. Simple linear regression analysis also showed a significant increase in the number of days in which *Alternaria* exceeded allergenic threshold (β = 1.82, R^2^ = 0.40, p = 0.02) while there was no significant trend in the number of days in which Cladosporium exceeded its allergenic threshold (β = −0.82, R^2^ = 0.03, p = ns) (Figure 4, Table S4). An analysis of season length showed only *Didymella* and *Tilletiopsis* to have a significantly increasing season length (*Didymella* β = 6.85, R^2^ = 0.47, p = 0.01, *Tilletiopsis* β = 4.51, R^2^ = 0.31, p = 0.05).

**Figure 4:**
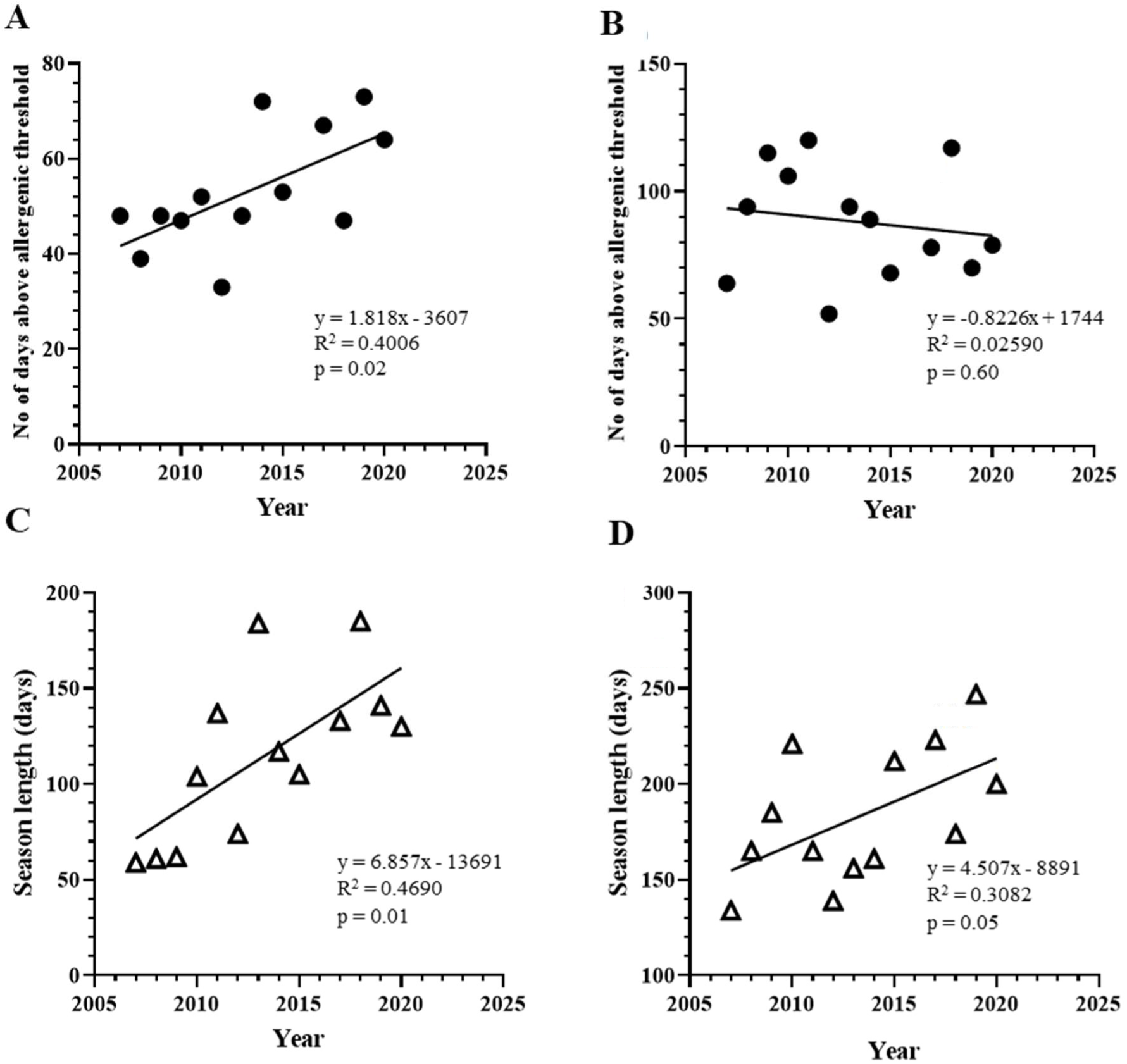
Linear regression analysis for annual changes in number of days above allergenic threshold (closed circles) for (**A**) *Alternaria* (> 100 spores/m^3^/day) and (**B**) *Cladosporium* (> 3000spores/m^3^/day) and season length (open triangles) for (**C**) *Didymella* and (**D**) *Tilletiopsis*

## 4 Discussion

This study aimed to highlight the spore distribution and seasonal patterns of fungal species found in central England, providing a fungal spore calendar that can be used to understand variabilities that may have important influences on agriculture and health. The impact of meteorological factors on spore levels were also considered. The fungal spore calendar concentrates on the 9 most prevalent fungal spores observed over the 13 years studied. Of these, three are known to cause human allergy – *Alternaria, Cladosporium* and *Aspergillus/Penicillium*, while *Leptosphaeria, Didymella,* coloured basidiospores*, Sporobolomyces, Tilletiopsis* and *Ustilago* have all been implicated as causal agents in cases of allergy and thunder storm asthma (Bush and Yunginger, 1987; Denning et al., 2006; Horner et al., 1998; Koivikko and Savolainen, 1988; Levetin et al., 2016). However, information on allergenic threshold concentrations is only available for *Alternaria* and *Cladosporium*.

### 4.1 Fungal spore seasons

Despite variability in fungal spore concentrations between years, spore distribution showed highly consistent patterns over the study period. The spore season in and around Leicester showed a bimodal trend, with the main peak period being in the summer months, which coincided with the greatest variety of spore species, followed by a smaller peak in autumn, mainly due to basidiospore release.

Spore concentrations consistently began to increase in May with *Ustilago*, a smut, being the first to reach its peak concentration in June, followed by *Cladosporium, Tilletiopsis and Sporobolomyces* whose concentrations increased throughout June, reaching maximal in July and August. *Alternaria* and *Didymella* levels were later to increase, rising in July and reaching their peak in August, while *Aspergillus/Penicillium* type and coloured basidiospore concentrations increased slowly during the summer, not reaching their peak until the autumn months. This annual distribution of spore species was consistent across the duration of this study despite wide variations in spore concentrations from year to year.

Comparable annual distribution patterns were observed in other studies, carried out at similar latitudes in Europe to Leicester (Corden and Millington, 2001; Martínez-Bracero et al., 2022; Nikkels et al., 1996; Sadys et al., 2016a). However, at locations with a more southerly latitude and Mediterranean climate these spores tended to appear earlier in the year and remain for a longer period of time (Anees-Hill et al., 2022; Antón et al., 2019; Katotomichelakis et al., 2016; Katsimpris et al., 2022; Reyes et al., 2016; Sousa et al., 2016). Of the more abundant spores observed in this study only *Leptosphaeria* showed a differing annual distribution compared to other sites. Derby and Worcester both reported a modest August peak for *Leptosphaeria* (Sadys et al., 2016a) while, Scevkova and Kovac reported a peak in July in a 1 year study in Bratislava (Scevkova and Kovac, 2019) and April in a 2 year study in central Spain (Reyes et al., 2016). In this study *Leptosphaeria* was not found to have a consistent, distinct season, peaking at different times in different years e.g., May in 2014, 2017 and 2018 and August in 2008 and 2012, explaining the difference between the current study and the data from the nearby Derby site. This clearly demonstrates the importance of longitudinal studies using data across a number of years when preparing a fungal spore calendar.

*Cladosporium* was the most abundant airborne fungal spore in this study, a feature found to be characteristic of a temperate climate (D’Amato and Spieksma, 1995) and observed in many studies across Europe (Antón et al., 2019; Kasprzyk et al., 2004; Martínez-Bracero et al., 2022; Sadys et al., 2016a; Scevkova and Kovac, 2019; Vélez-Pereira et al., 2021).

However, SSIn and/or maximal daily peak concentrations of Cladosporium were found to be several fold higher in Leicester than in many of these studies. Distance from the sea, proximity to agricultural land and /or green spaces as well as a mean temperature above 16°C and relative humidity of 80% have all been shown to favour *Cladosporium* spore levels (Kasprzyk et al., 2021; Katial et al., 1997; Sindt et al., 2016) and were all applicable to the Leicester area during the summer months, possibly contributing to the high spore concentrations.

The concentrations of the highly allergenic *Alternaria* spores were also found to be significantly higher in this study than in most other investigations across Europe. Only studies from Bratislava, Slovakia (Scevkova and Kovac, 2019), Saclay, France (Roland Sarda-Estève et al., 2019) and Lleida, Spain (Vélez-Pereira et al., 2016) showed similar or higher concentrations. Despite a range of geographical locations all share a common feature with Leicester – a rural location or proximity to agricultural land. Apangu et al. (2020 showed that substantially higher concentrations of *Alternaria* spores were associated with areas with high amounts of cereals and oilseed rape and peak periods of spore concentration tend to coincide with harvest time (Corden and Millington, 2001).

The spikes in spore levels observed for *Cladosporium* (July) and *Alternaria* (August) alongside their high maximal daily peaks could have important implications for those with fungal atopy. Information presented in this study could provide important information to inform effective symptom management strategies.

### 4.2 Spore Dependence on Meteorological Parameters

Concentrations of the “dry weather spores”, *Alternaria* and *Cladosporium*, were significantly positively correlated to daily temperature. *Alternaria* concentrations showed a negative relationship with precipitation, however multivariate analysis indicated that *Cladosporium* had a positive correlation with precipitation on the previous day. These differences in the effects of meteorological parameters on *Alternaria* and *Cladosporium* may reflect differing requirements for sporulation of the two taxa, *Cladosporium* requiring a lower temperature (13 – 21 °C) and more humid environment (Katial et al., 1997; Sabariego et al., 2000). *Alternaria’s* higher optimal growing temperature (22 – 28 °C) (Fernández et al., 1998) corresponds with the later seasonal appearance and shorter season observed for *Alternaria*.

The variance in *Ustilago* spore concentrations was influenced by maximum daily temperature and precipitation on the previous day, however these meteorological parameters had a smaller predictive value for *Ustilago* than for *Cladosporium*. This along with the consistent, short season suggests other factors, such as the availability of a suitable host, plays a role in the presence of *Ustilago* spores. The June peak for *Ustilago* reported in this study corresponds to the peak in grass pollen in the Leicester area, this aligns with the data from Crotzer and Levetin (1996 who found that *Ustilago* peaks in May and June in Tulsa, Oklahoma in conjunction with the flowering seasons of wheat, oats and other grasses.

The presence of the “wet weather spores” of *Didymella*, *Sporobolomyces*, *Tilletiopsis* and *Leptosphaeria* all correlated with precipitation in agreement with studies from Dublin (Martínez-Bracero et al., 2022), Worcester (Sadys et al., 2016a) Bratislava (Scevkova and Kovac, 2019) and Salamanca, Spain (Antón et al., 2019). However, the meteorological parameters used in the multiple regression analysis only accounted for 10 – 20% of the variance in the spores of these species. Other studies have indicated that relative humidity, leaf wetness, dew point temperature and minimum air temperature can be more important factors in relation to “wet weather spore” concentrations (Richardson, 1996; Sadys et al., 2018; Sadys et al., 2016b; Stępalska et al., 2012) and as such may be more appropriate indicators of “wet weather spore” concentrations than total precipitation.

The main meteorological factors influencing fungal spore concentration are temperature and precipitation as seen in this study and summarised by Anees-Hill et al. (2022 in a survey of studies from around Europe. These parameters also provide optimal growth conditions for plants which form the predominant growth material for fungal organisms. Thus, the peak in spore concentrations during summer and early autumn months observed in this study and others may in part be due to favourable climatic and weather conditions allowing the growth and/or decomposition of plant material as well as the growth and spore release of the fungi itself.

### 4.3 Changes in seasonal fungal spore concentration

The long-term trend for total spore SSIn, over the 13 years of this study, was a statistically significant decrease, however this decrease appears to result from decreases in the concentrations of the “wet weather spores” of *Sporobolomyces* and *Tilletiopsis,* which reached significance, and the decreasing trend for *Didymella* and *Leptosphaeria*, which together make up 4 of the top 5 most abundant spores, identified to genus level, in the air around Leicester. This decreasing trend for “wet weather spore” concentrations coincided with a significant decrease in daily precipitation.

Few studies have looked at the long term trend for “wet weather spores” despite their potential as allergens and links to thunderstorm induced asthma (Denning et al., 2006). However, of those that have, a decreasing trend has been noted in the number of “wet weather spores” including; ascospores and basidiospores in Saclay, France (Sarda-Estève et al., 2019), *Leptosphaeria* spore concentrations, Thessaloniki, Greece (Damialis et al. (2015 and Catalonia, Spain (Vélez-Pereira et al., 2016) and, and *Didymella,* Worcester, UK (Sadys et al., 2016a).

While the “wet weather spores” showed a decreasing trend in this study, the highly allergenic “dry weather spores” of *Alternaria* and *Cladosporium* showed an increasing trend which reached statistical significance for *Alternaria*. It was also found that the number of days in which *Alternaria* exceeded allergenic threshold each year demonstrated a significant increase. It has been speculated that changes in local land use over time may have important impacts on *Alternaria* concentrations. Studies from Cardiff, Copenhagen, Bratislava and Bellaterra (NE Spain) have shown a decline in agricultural use and the growth of cereal crops was a driving force for the decline in *Alternaria* concentrations (Corden et al., 2003; Olsen et al., 2020; Scevkova et al., 2016; Vélez-Pereira et al., 2016). Leicester is surrounded by agricultural land and has a high proportion of cereal crops and oil seed rape, major hosts for *Alternaria* (Apangu et al., 2020). The peak for *Alternaria* spore concentrations, in August coincides with peak cereal harvesting, an activity long associated with high emissions of *Alternaria* spores. Thus, the increase in *Alternaria* spores in this study may be linked to increasing agricultural activity around Leicester, as no significant link to increasing temperature was observed.

The widespread distribution of *Cladosporium* and its dominance over other fungal airspora, irrespective of urban or rural location, suggests levels of *Cladosporium* spores are less dependent on local agricultural trends so did not show the same increase over time as *Alternaria* in this study. The multiple regression analysis of meteorological variables on fungal spore concentrations here also indicated that daily precipitation had a negative impact on *Alternaria* spore concentrations but a positive effect on *Cladosporium* when a lag of 1 day was considered, in agreement with Sadys (2017. Nikkels et al. (1996 noted that *Cladosporium* spore concentrations were “extraordinarily high” during a very hot and humid summer. Although no change in average temperature was observed across the duration of this study a decreasing trend for daily precipitation was indicated, which may have influenced the rise in *Alternaria* spores and had a negative impact on *Cladosporium*.

### 4.4 Strengths and Limitations

This study has the advantage of being one of the longest sampling studies of fungal spores found in the outdoor air, and the longest multi-taxa study in the UK, which is important in addressing the year-to-year variability in fungal spore concentrations and distribution. It also looks at a wide range of spores from both the “dry” and “wet” weather airspora and relates their concentrations to meteorological parameters. A limitation is that allergenic thresholds have only been established for *Cladosporium* (> 3000 spores/m^3^ per day) and *Alternaria* (> 100 spores/m^3^ per day) (Bagni et al., 1977; Gravesen, 1979; Rapiejko et al., 2007). More research into thresholds for other types of spores would help inform clinical advice with respect to spore counts. This study is based on the results from only one geographical location and as shown by Sindt et al. (2016), Vélez-Pereira et al. (2016), Anees-Hill et al. (2022) and Corden et al. (2003), distance from the coast, latitude, altitude and vegetation and agricultural activity in the vicinity of the sample site can all impact on the concentration and diversity of fungal spores observed and on long term trends in spore levels. However, Leicester is approximately in the centre of England, and as such should serve as a reasonable guide for airborne spores for central England and potentially wider areas of England and the UK. This study used a limited range of meteorological parameters but others that could have been considered were relative humidity, leaf wetness and dew point temperature which have all been shown to correlate closely with the concentrations of certain fungal spores. Finally, we did not examine the sources of spores – an analysis of the surrounding region’s vegetation and agricultural practices and changes in land use would potentially provide insights into changing trends in spore concentrations.

## 5 Conclusions

This study is amongst the longest sampling studies of fungal spores found in the outdoor air and the longest multi-taxa study in the UK. It provides a calendar for the seasonal distribution of the predominant spores found in the air around Leicester, UK in central England, which can help allergists determine the type of fungi patients may be sensitised to and aid disease management.

There is a growing appreciation of the impact of fungal spores on health and the economy and there is increasing evidence to suggest that climate change will impact aeroallergen levels (Lam et al., 2024). As demonstrated here, long term aeroallergen monitoring sites are crucial to investigating the effect of climate change on airborne fungal spores, and to understand the associated health implications.

## Acknowledgements

The authors would like to thank Awab Amin who carried out initial analysis

## Abbreviations

MSS: main spore season
P: precipitation
RMSE: root mean square error
SSIn: seasonal spore integral
Tm: maximum air temperature
WD: wind direction
WS: wind speed

## Supplementary Data

**Table S1.** Number of days sampled (maximum 275) and dates of missing spore counts. Missing dates were due to unforeseen circumstances including trap malfunctions.

| Year | Missing Dates | No of days<br>sampled |
| --- | --- | --- |
| 2007 | 01/04, 17-19/06, 21/06, 10/07, 3/09, 5-6/09, 11/09, 31/10, 19-27/11, 30/11 | 254 |
| 2008 | 01/11, 26/11, 30/11 | 272 |
| 2009 |  | 275 |
| 2010 |  | 275 |
| 2011 |  | 275 |
| 2012 |  | 275 |
| 2013 |  | 275 |
| 2014 | 22-23/06, 27-27/06, 2/07, 9-10/07, 26-28/07, 30-31/07 | 263 |
| 2015 | 18-31/03, 3/11 | 260 |
| 2017 | 29-31/03 | 272 |
| 2018 | 2-9/10, 4-31/11 | 240 |
| 2019 |  | 275 |
| 2020 | 19-23/06, 5/10, 19-26/11 | 261 |

**Table S2.**
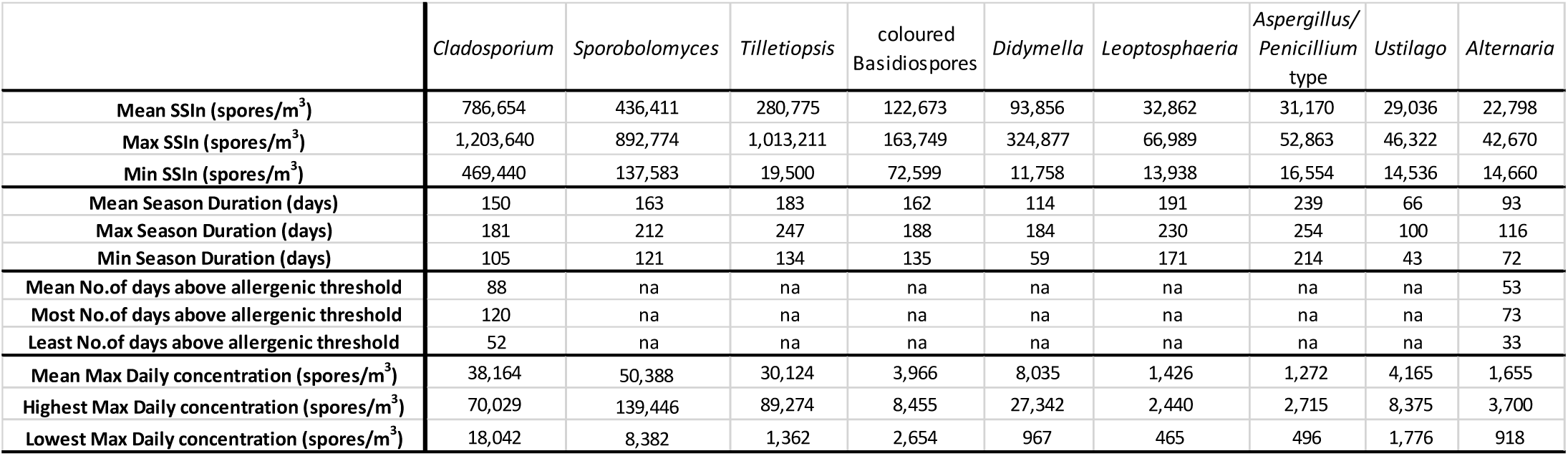
Seasonal characteristics of the main spore types studied

|  | *Cladosporium* | *Sporobolomyces* | *Tilletiopsis* | coloured<br>Basidiospores | *Didymella* | *Leptosphaeria* | *Aspergillus/*<br>*Penicillium*<br>type | *Ustilago* | *Alternaria* |
| --- | --- | --- | --- | --- | --- | --- | --- | --- | --- |
| Mean SSIn (spores/m<sup>3</sup>) | 786,654 | 436,411 | 280,775 | 122,673 | 93,856 | 32,862 | 31,170 | 29,036 | 22,798 |
| Max SSIn (spores/m<sup>3</sup>) | 1,203,640 | 892,774 | 1,013,211 | 163,749 | 324,877 | 66,989 | 52,863 | 46,322 | 42,670 |
| Min SSIn (spores/m<sup>3</sup>) | 469,440 | 137,583 | 19,500 | 72,599 | 11,758 | 13,938 | 16,554 | 14,536 | 14,660 |
| Mean Season Duration (days) | 150 | 163 | 183 | 162 | 114 | 191 | 239 | 66 | 93 |
| Max Season Duration (days) | 181 | 212 | 247 | 188 | 184 | 230 | 254 | 100 | 116 |
| Min Season Duration (days) | 105 | 121 | 134 | 135 | 59 | 171 | 214 | 43 | 72 |
| Mean No.of days above allergenic threshold | 88 | na | na | na | na | na | na | na | 53 |
| Most No.of days above allergenic threshold | 120 | na | na | na | na | na | na | na | 73 |
| Least No.of days above allergenic threshold | 52 | na | na | na | na | na | na | na | 33 |
| Mean Max Daily concentration (spores/m<sup>3</sup>) | 38,164 | 50,388 | 30,124 | 3,966 | 8,035 | 1,426 | 1,272 | 4,165 | 1,655 |
| Highest Max Daily concentration (spores/m<sup>3</sup>) | 70,029 | 139,446 | 89,274 | 8,455 | 27,342 | 2,440 | 2,715 | 8,375 | 3,700 |
| Lowest Max Daily concentration (spores/m<sup>3</sup>) | 18,042 | 8,382 | 1,362 | 2,654 | 967 | 465 | 496 | 1,776 | 918 |

**Table S3.**
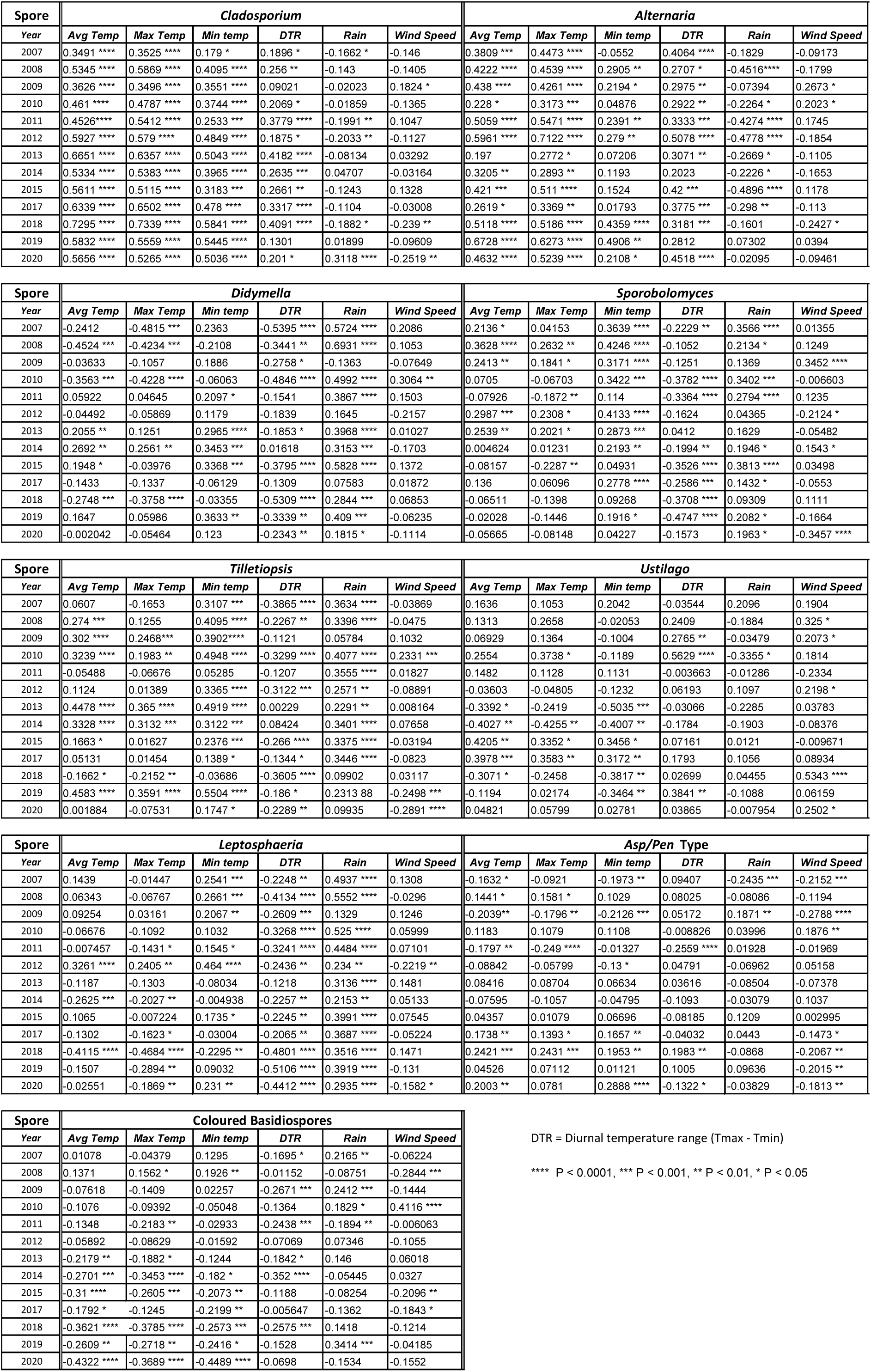
Spearman’s rank correlations between daily spore concentration and measured weather parameters during the spore season.

**Table S4.** Linear regression analysis of SSIn and year for each fungal spore type counted

|  | Slope | R<sup>2</sup> | Deviation from zero |
| --- | --- | --- | --- |
| *Cladosporium* | 5751 ± 14866 | 0.01342 | ns |
| *Alternaria* | 1352 ± 505.2 | 0.3944 | P = 0.02 |
| *Didymella* | -11365 ± 6014 | 0.2451 | P = 0.09 |
| *Sporobolomyces* | -32819 ± 13137 | 0.3620 | P = 0.03 |
| *Tilletiopsis* | -37315 ± 16506 | 0.3172 | P = 0.05 |
| *Ustilago* | -362.1 ± 656.9 | 0.02688 | ns |
| *Leptosphaeria* | -1741 ± 1020 | 0.2093 | ns |
| *Asp/Pen* type | 465.9 ± 600.1 | 0.05195 | ns |
| Coloured basidiospores | 616 ± 2036 | 0.008254 | ns |
| Total spores | -105641 ± 42813 | 0.3563 | P = 0.03 |

SSIn – seasonal spore interval (i.e., total spore concentration during the spore season as
defined by the 90% method)

